# A role of the Nse4 kleisin and Nse1/Nse3 KITE subunits in the ATPase cycle of SMC5/6

**DOI:** 10.1101/760678

**Authors:** L. Vondrova, M. Adamus, M. Nociar, P. Kolesar, A.W. Oliver, J.J. Palecek

## Abstract

The SMC (Structural Maintenance of Chromosomes) complexes are composed of SMC dimers, kleisin and kleisin-interacting subunits. Mutual interactions of these subunits constitute the basal architecture of the SMC complexes. Particularly, terminal domains of the kleisin subunit bridge the SMC head domains of the SMC molecules. Binding of ATP molecules to the heads and their hydrolysis alter the shape of long SMC molecules (from rod-like to open-ring shapes) and power them with motor activity. The kleisin-interacting (HAWK or KITE) subunits bind kleisin linker regions and regulate such dynamic activities.

We developed new systems to follow the interactions between SMC5/6 subunits and stability of the SMC5/6 complexes. First, we show that the N-terminal domain of the Nse4 kleisin molecule binds to the SMC6 neck and bridges it to the SMC5 head. Second, binding of the Nse1 and Nse3 KITE proteins to the Nse4 linker part increases stability of the ATP-free SMC5/6 complex. In contrast, binding of ATP to the SMC5/6 complex containing KITE subunits significantly decreases its stability. Elongation of the Nse4 linker partially suppresses instability of the ATP-bound complex, suggesting that the binding of the KITE proteins to the Nse4 linker constrains its limited size. Our data suggest that the KITE proteins may shape the Nse4 linker to fit the ATP-free complex optimally and to facilitate opening of the complex upon ATP binding. This mechanism suggests an important role of the KITE subunits in the dynamics of the SMC5/6 complexes.

**AUTHOR SUMMARY:** The SMC5/6 complex is member of the Structural maintenance of chromosomes (SMC) family, key organizers of both prokaryotic and eukaryotic genomes. Their architecture and dynamics (driven by ATP binding and hydrolysis) are essential for cellular processes, like chromatin segregation, condensation, replication and repair. In this paper, we described conserved mode of the Nse4 kleisin subunit binding to the SMC6 (similar to cohesin and condensin) and its bridging role. Furthermore, we showed different impact of the binding of the Nse1-Nse3 KITE subunits to the Nse4 kleisin bridge. Our study suggested that the KITE proteins modulate the stability of the SMC5/6 complex, depending on its binding and hydrolysis of ATP. Our findings uncover molecular mechanisms underlying dynamics of the SMC5/6 complexes.

## INTRODUCTION

The SMC (Structural Maintenance of Chromosomes) complexes are key organizers of prokaryotic and eukaryotic genomes. They organize chromatin domains (cohesins; [1]), condense mitotic chromosomes (condensins; [2]), assist in DNA repair (SMC5/6; [3, 4]) and replication (SMC/ScpAB; [5]). These circular complexes use the energy of ATP hydrolysis to drive DNA topology changes. In prokaryotes, SMC/ScpAB drives extrusion of loops forming behind the replication fork. In eukaryotes, condensins extrude loops laterally and axially to shape chromatin to the typical mitotic chromosomes. Cohesins assist in formation of topologically associating domains during interphase. Cohesin rings can also hold newly replicated sister chromatids together and release them in highly controlled manner. The SMC5/6 complexes have been implicated in the repair of DNA damage by homologous recombination, stabilization and restart of stressed replication forks. The SMC5/6 instability leads to the chromosome breakage syndrome in human [6], however, the molecular mechanism of the SMC5/6 action is largely unclear.

All the SMC complexes are composed of three common categories of subunits: SMC, kleisin and kleisin-interacting proteins [7, 8]. The SMC proteins are primarily build of long anti-parallel coiled-coil arms, a globular hinge (situated in the middle of their peptide chain) and a head domain (formed by combined amino and carboxyl termini; [9–13]). The globular head domain contains ATP binding and hydrolysis motifs of the ATP-binding cassette transporter family [14, 15]. Two SMC molecules form dimers via the association of their hinge domains and transiently interact when their head domains sandwich a pair of ATP molecules. The binding of ATP changes conformation and shape of the SMC subunits at local as well as global levels [10]. At local level, the SMC heads and necks move from aligned position to the ATP-locked conformation. At global level, the overall shape of the complex changes from rod- to ring-like upon ATP binding (with heads locked by ATP at one end and hinge dimer at the other end). The hydrolysis of ATP dissolves the SMC-ATP-SMC head bridge.

The ATPase head domains are also connected by the kleisin subunit in an asymmetric way. Kleisin binds to the cap side of one SMC (designated as κSMC) head domain via a winged-helix domain (WHD) at its carboxyl terminus. Kleisin’s α-helix located at its amino terminal helix-turn-helix (HTH) domain binds to the coiled-coil base region immediately adjacent to the other SMC head (called neck and designated as νSMC; [16–18]). This kleisin bridge mediated by protein-protein interactions seems to be more permanent than the ATP-mediated bridge (as the latter bridge dissolves upon ATP hydrolysis). However, the ATP binding may induce dissociation of the kleisin-νSMC interaction and lead to release of DNA from cohesin ring [19–24].

The kleisin-interacting subunits bind and shape kleisin linker regions conecting N-(HTH) and C-terminal (WHD) domains. The KITE (Kleisin-Interacting Tandem winged-helix Element) subunits interact with kleisins in prokaryotic SMC/ScpAB and eukaryotic SMC5/6 complexes, while eukaryotic cohesin and condensin complexes associate with HAWK (HEAT proteins Associated With Kleisin) proteins. Interestingly, HAWK (Scc3 and Pds5) and Wapl proteins regulate release of cohesin from chromosomes. It was proposed that these proteins shape and stiffen the linker region of kleisin molecule, which assists in transduction of conformational changes of the ATP-mediated SMC head dimerization to the dissociation of the kleisin-νSMC interaction (Scc1-Smc3 in the cohesin complex). Similarly, KITE proteins assist in opening of the SMC/ScpAB complex [25, 26]. However, the role of the KITE subunits in the SMC5/6 complex remains largely elusive.

Here we aimed to uncover relationships between SMC5/6 subunits and their roles in the SMC5/6 dynamics. We developed new systems to analyse the interactions between SMC5/6 subunits and to follow the stability of SMC5/6 complexes. Using systems composed of SMC6-Nse4-Nse3-Nse1 and SMC6-Nse4-SMC5, respectively, we showed that the N-terminal HTH domain of the Nse4 kleisin molecule binds to the SMC6 neck and bridges it with the SMC5 head. With a more complex system, we observed increased stability of the ATP-free SMC5/6 complex upon binding of the Nse1 and Nse3 KITE proteins to the Nse4 kleisin. While the ATP-free complex was highly stable, the binding of ATP to the SMC5/6 complex containing KITE-bound kleisin significantly decreased its stability. Reduced Nse3 binding to the Nse4 linker or elongation of the Nse4 linker partially suppressed the instability of the ATP-bound complex, suggesting that the binding of the KITE proteins to the Nse4 linker constrains its limited size. Our data suggest that the KITE proteins may shape the Nse4 linker to fit the ATP-free complex optimally and to facilitate opening of the complex upon ATP binding.

## RESULTS

### Nse4 interacts with SMC6 neck

We have previously shown that Nse4 belongs to the kleisin superfamily of proteins and binds to SMC5 and SMC6 head fragments [27]. To map Nse4 interactions with SMC6, we employed peptide libraries covering yeast *S. pombe* SMC6 regions aa 241-365 and aa875-1024 (Suppl. Fig. S1A and Supplementary Table 1; unpublished data). The peptides were pre-bound to ELISA plates and tested against Nse4(1-150) and control (human TRF2) protein [28, 29]. The peptides covering the C-terminal region of SMC6 (aa955-1009) bound to Nse4 (Figs. 1A and S1A). The peptide aa960-984 exhibited the highest affinity and specificity to Nse4 while other peptides bound to Nse4 in a less specific way (e.g. peptide aa970-994). Interestingly, the SMC6 region aa960-984 corresponds to the SMC neck regions interacting with kleisins in most SMC complexes [12].

To analyse the Nse4-SMC6 interaction in more detail, we established various multicomponent yeast two-hybrid (mY2H) systems [30]. It was difficult to follow the Nse4-SMC6 binary interaction in classical Y2H (Fig. 1B, column 3; [27, 31, 32]), therefore we added DNA encoding Nse1 and Nse3 subunits on an extra plasmid (p416ADH1-Nse1+Nse3 construct, 4Y2H) to enhance Nse4-binding properties [33]. Indeed, addition of both Nse1 and Nse3 subunits to Gal4BD-Nse4/Gal4AD-SMC6 resulted in a stable SMC6-Nse4-Nse3-Nse1 complex (Fig. 1B, column 4). Similarly, addition of SMC5 to Nse4-SMC6 (3Y2H) resulted in formation of a stable SMC5-Nse4-SMC6 complex (Fig. 1C, column 4).

**Figure 1.**
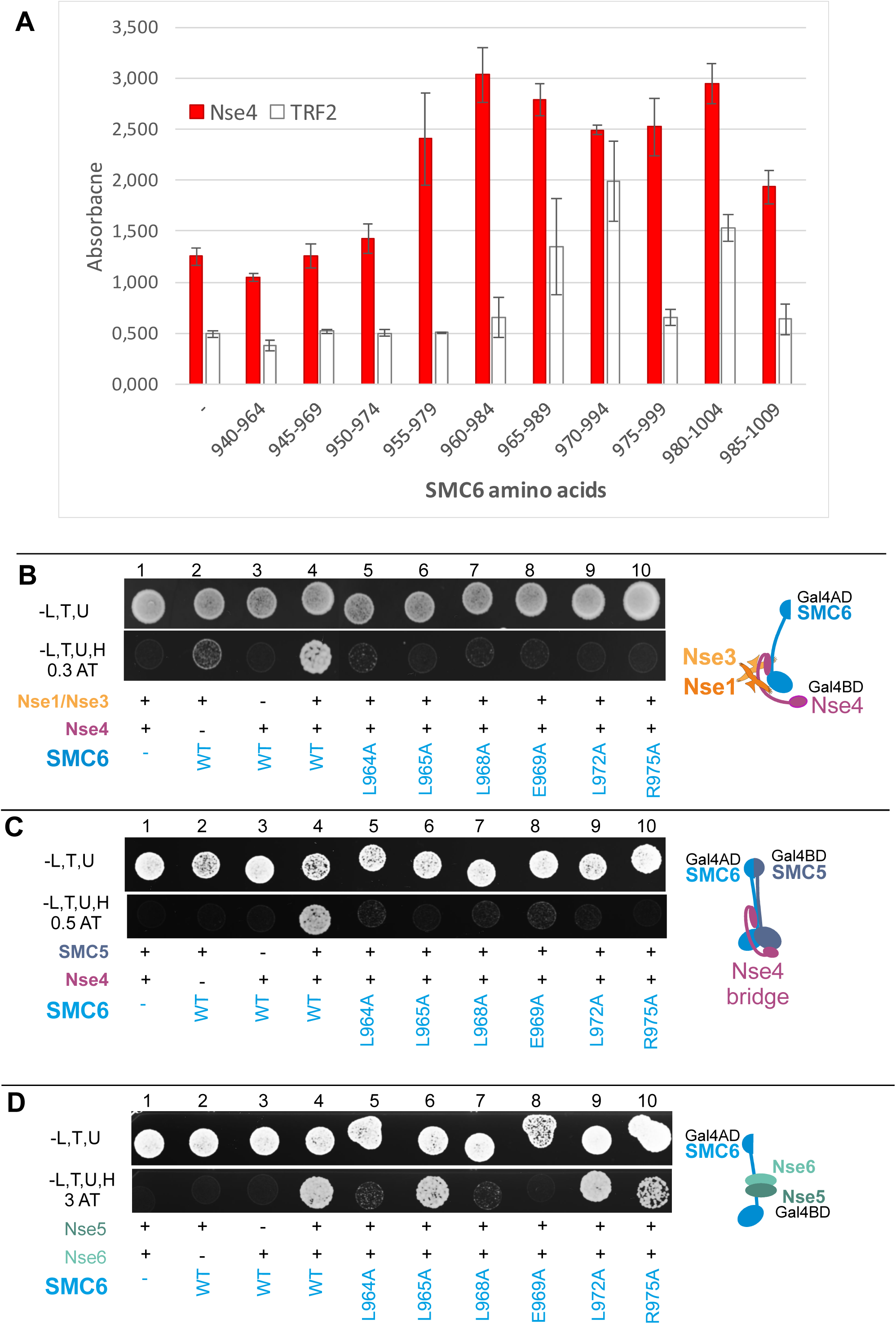
Nse4 binds neck region of the SMC6 protein. Peptide library (A) and multi-component yeast two-hybrid systems (B-D) were employed to determine SMC6 region and residues binding to Nse4. (A) Quantification of relative binding of the Nse4(1-150) protein (Nse4; red columns) to the SMC6 synthetic peptides (listed in Suppl. Table S1) using the PEPSCAN-ELISA method. The SMC6(aa960-984) peptide exhibits the highest affinity and specificity to Nse4. Results show mean ± SEM of 3 independent measurements. His-TRF2 protein (TRF2; white columns) was used in the control experiment. (B-D) Impact of mutations on the stability of the following SMC6 complexes was tested: SMC6-Nse4-Nse3-Nse1 (B), SMC6-Nse4-SMC5 (C) and SMC6-Nse5-Nse6 (D). Schematic representation of the complexes is at the right side of each panel. (B) The full-length hybrid SMC6 (fused to Gal4AD domain) and Nse4 (fused to Gal4BD domain) constructs were co-transformed together with Nse1+Nse3 plasmid (p416ADH1 vector backbone) into *S. cerevisiae* PJ69 cells. Formation and stability of the SMC6-Nse4-Nse3-Nse1 complex was scored by growth of yeast PJ69 transformants on plates without leucine, tryptophan, uracil and histidine, containing 0.3 mM 3-Amino-1,2,4-triazole (-L,T,U,H, 0.3AT panel). The L964A, L965A, L968A, E969A, L972A and R975A mutations reduce stability of the SMC6 complexes. (C) Similarly, the full-length hybrid SMC6 (fused to Gal4AD domain) and SMC5 (fused to Gal4BD domain) were co-transformed together with non-hybrid Nse4 full-length construct (p416ADH1 vector) and stability of the SMC5-Nse4-SMC6 complex was scored on plates containing 0.5 mM 3-Amino-1,2,4-triazole (-L,T,U,H, 0.5AT panel). The L964A, L965A, L968A, E969A, L972A and R975A mutations reduce stability of the SMC6 complexes. (D) In the control experiment, the same mutations were tested in the SMC6-Nse5-Nse6 complex (constituted of the full-length Gal4AD-SMC6, Gal4BD-Nse5 and non-hybrid Nse6). Stability of the SMC6-Nse5-Nse6 complex was scored on plates containing 3 mM 3-Amino-1,2,4-triazole (-L,T,U,H, 3AT panel). The L964A, L968A and E969A mutations affect all SMC6 complexes (B-D) while L965A, L972A and R975A mutations reduce only stability of SMC6-Nse4 complexes (B and C), suggesting that the highly conserved L965, L972 and L975 residues (Suppl. Fig. S1B) are specifically required for the SMC6 interaction with Nse4. Wild-type (WT) or mutant versions of SMC6 are labelled in blue below the panels; “-“, control empty vector; “+”, co-transformed construct (as labelled at the left side). Growth of the transformants was verified on control plates without leucine, tryptophan and uracil (-L,T,U). All mY2H tests were repeated at least 3 times.

Using these mY2H systems and site-directed mutagenesis, we aimed to identify the Nse4-binding residues within the most conserved part of the ELISA-defined SMC6 region (aa960-984; Figs. 1A and S1B; unpublished data). The L964A, L965A, L968A, E969A, L972A and R975A mutations reduced stability of the SMC6-Nse4-Nse3-Nse1 tetramer while the others had negligible effect (Figs. 1B and S1C). These data suggest that residues L964, L965, L968, E969, L972 and R975 may mediate either the Nse4-SMC6 interaction or putative interactions between SMC6 and Nse1-Nse3 subunits. To exclude the latter possibility, we employed 3Y2H system consisting of the SMC5/SMC6/Nse4 subunits. Again, L964A, L965A, L968A, E969A, L972A and R975A mutations reduced stability of the (SMC5-)SMC6-Nse4 complex while the others had no effect (Figs. 1C and S1D), suggesting that these residues may mediate Nse4-SMC6 interaction.

To distinguish between mutations specifically disturbing the Nse4-SMC6 interaction from those affecting SMC6 structure, we established another 3Y2H system consisting of SMC6 and Nse5-Nse6 subunits (as Nse5 and Nse6 bind to the SMC6 protein; [27]). In this system, we used the same Gal4AD-SMC6 mutation constructs as above in combination with Gal4BD-Nse5 and p416ADH1-Nse6 (Figs. 1D and S1E). The mutations L964A, L968A and E969A reduced SMC6-Nse5-Nse6 complex stability, suggesting their deleterious effect on SMC6 structure (Fig. 1, compare panels B, C and D, columns 5, 7 and 8). In contrast, the other mutations had no impact, suggesting that the conserved L965, L972 and R975 residues within the SMC6 neck region mediate SMC6-Nse4 interaction (Figs. 1 and S1).

### SMC6 binds N-terminal motif of Nse4 protein

Next, we took advantage of the crystal structures of the kleisin-νSMC complexes [16–18, 26, 34], which suggest a key role for the N-terminal HTH domain of the kleisin molecule in its binding to νSMC neck. We mutated residues within the third α-helix of the Nse4 HTH domain (aa62-68) and analysed their impact on the Nse4-SMC6 interaction using the SMC5-Nse4-SMC6 3Y2H system as above (Fig. 1C). The Nse4 mutations L62C, K64C, T65R, D67C and L68C reduced the SMC5-Nse4-SMC6 stability significantly while the others had no effect (Fig. 2A). To distinguish between mutations specifically disturbing Nse4-SMC6 interactions and mutations affecting Nse4 structure, we tested the Gal4BD-Nse4 mutation constructs in combination with Gal4AD-Nse3 in the classical Y2H system (Fig. 2B; [28, 35]). The K64C and D67C mutations affected the Nse3-Nse4 interaction, suggesting their deleterious effect on the Nse4 structure (Fig. 2B, columns 6 and 9). In contrast, the other mutations had no effect on the Nse3-Nse4 interaction, suggesting that the intact L62, T65 and L68 conserved residues are required for the Nse4-SMC6 binding (Fig. 2C). Altogether, our data suggest that the Nse4 HTH motif binds the SMC6 neck region and that the SMC6-Nse4 interaction mode is similar to the other νSMC-kleisin interactions [12, 13].

**Figure 2.**
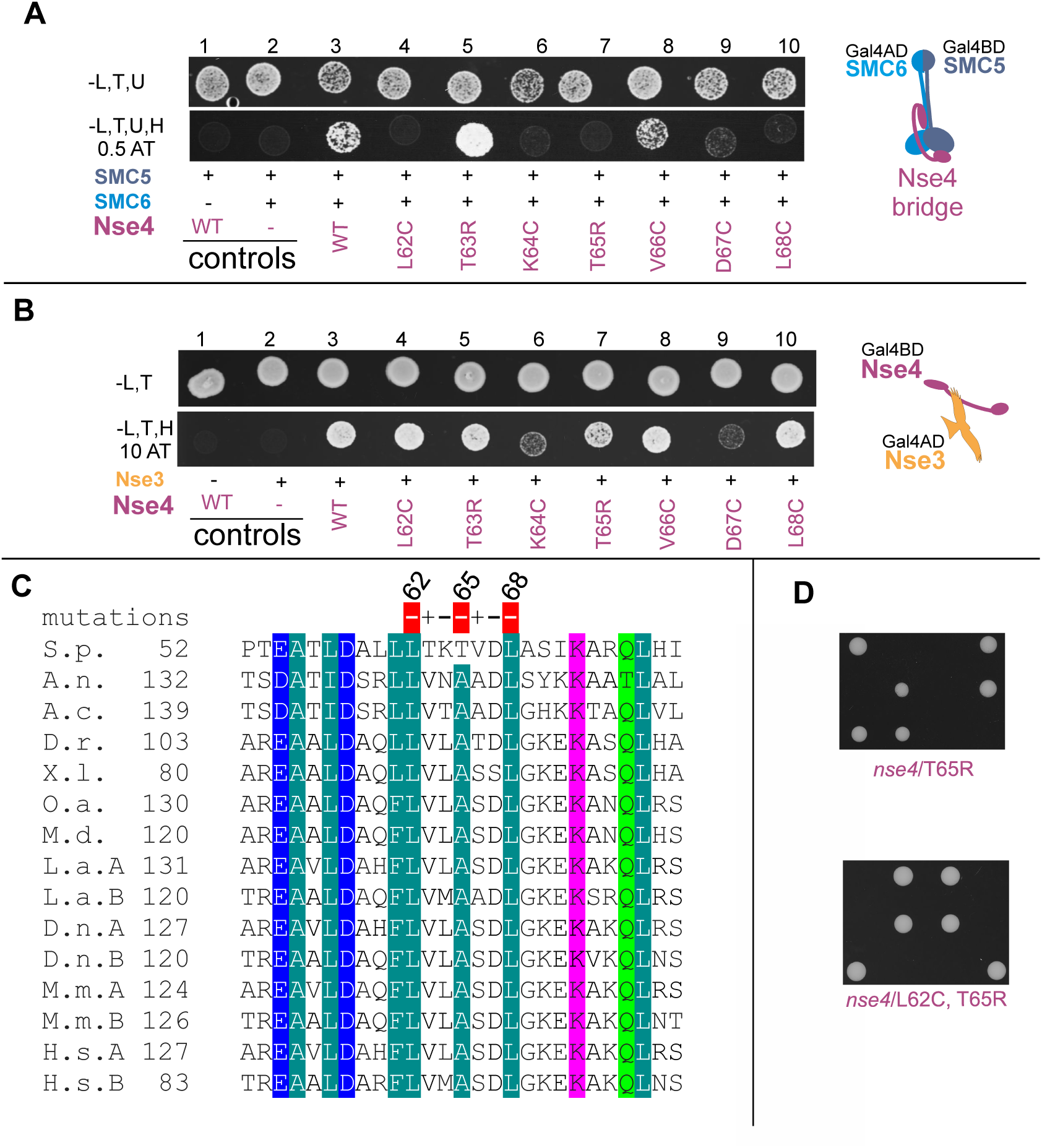
Binding of the Nse4 helix H3 to SMC6 is essential for yeast viability. Identification of the SMC6-binding residues within the Nse4 helix H3 region (aa62-68). (A) Stability of the SMC5-Nse4-SMC6 complex was scored by 3Y2H on plates containing 0.5 mM 3-Amino-1,2,4-triazole (-L,T,U,H, 0.5 AT panel; further details as in Fig. 1). The Nse4/L62C, K64C, T65R, D67C, and L68C mutations reduce stability of the SMC6 complexes. In the control experiment (B), the same mutations were tested for the Nse4-Nse3 interaction in the classical Y2H system (constituted of the full-length Gal4AD-Nse4 and Gal4BD-Nse3). Interactions were scored on plates containing 10 mM 3-Amino-1,2,4-triazole (-L,T,U,H, 10AT panel). The K64C and D67C mutations affect all Nse4 complexes (A and B). Intact L62, T65 and L68 Nse4 residues are required specifically for binding to SMC6. Wild-type (WT) or mutant versions of Nse4 are labelled in violet below the panels (further details as in Fig. 1). (C) Alignment of the Nse4 helix H3 (of the N-terminal HTH domain). The orthologs are from *Schizosaccharomyces pombe* (*S.p.*), *Aspergillus nidulans* (*A.n.*), *Aspergillus clavatus* (*A.c.*), *Danio rerio* (*D.r.*), *Xenopus laevis* (*X.l.*), *Ornithorhynchus anatinus* (*O.a.*), *Monodelphis domestica* (*M.d.*), *Loxodonta africana* (*L.a.*), *Dasypus novemcinctus* (*D.n.*), *Mus musculus* (*M.m.*), *Homo sapiens* (*H.s.*). Note that there are two Nse4 genes in the placental mammals denoted as A and B. “+”, mutation not affecting Nse4 interactions; “-“, mutation disrupting all Nse4 complexes; red minus, mutation specifically disrupting the Nse4-SMC6 interaction. Amino acid shading represents following conserved amino acids: *dark green*, hydrophobic and aromatic; *light green*, polar; *pink*, basic; *blue*, acidic. (D) Tetrad dissection analysis of yeast *S. pombe* diploid strains *nse4^+^/nse4-T65R* (top) and *nse4^+^/nse4-L62C, T65R* (bottom). The *nse4-T65RC* and *nse4-L62C, T65R* mutations are lethal, suggesting essential role of the Nse4–SMC6 interaction.

To analyse the role of the Nse4-SMC6 interaction in yeast cells, we introduced the L62C and T65R mutations into the genome of diploid *S. pombe*. Tetrad analysis showed that the single *nse4*-*T65R* and double *nse4*-*L62C, T65R* mutations were lethal (Fig. 2D), suggesting an essential role for the Nse4-SMC6 interaction.

### A role for the Nse4 and ATP molecules in bridging of SMC5-SMC6

To compare the role of Nse4 and ATP in bridging of the SMC5-SMC6 heads, we introduced the SMC5/E995Q mutation which inhibits ATP hydrolysis; i.e. enhances ATP retention between SMC5-SMC6 heads and their dimerization (Figs. 3A and S2; [14, 36]). In the 3Y2H system, the interaction between Gal4BD-SMC5 and Gal4AD-SMC6 constructs was not detectable, suggesting a low stability of the SMC5-SMC6 dimer even upon stable binding of ATP (Fig. 3A, columns 1 and 2). Addition of Nse4 resulted in stable SMC5-Nse4-SMC6 complex formation (Fig. 3A, columns 3 and 4), suggesting that Nse4 stabilizes the bridge between SMC5 and SMC6. The introduction of the ATP-hydrolysis mutation to the SMC5-Nse4-SMC6 complex only slightly increased its stability (Fig. 3A), suggesting a major role of Nse4 (and a minor additive effect of ATP binding; see below) in bridging of the SMC5-SMC6 heads.

**Figure 3.**
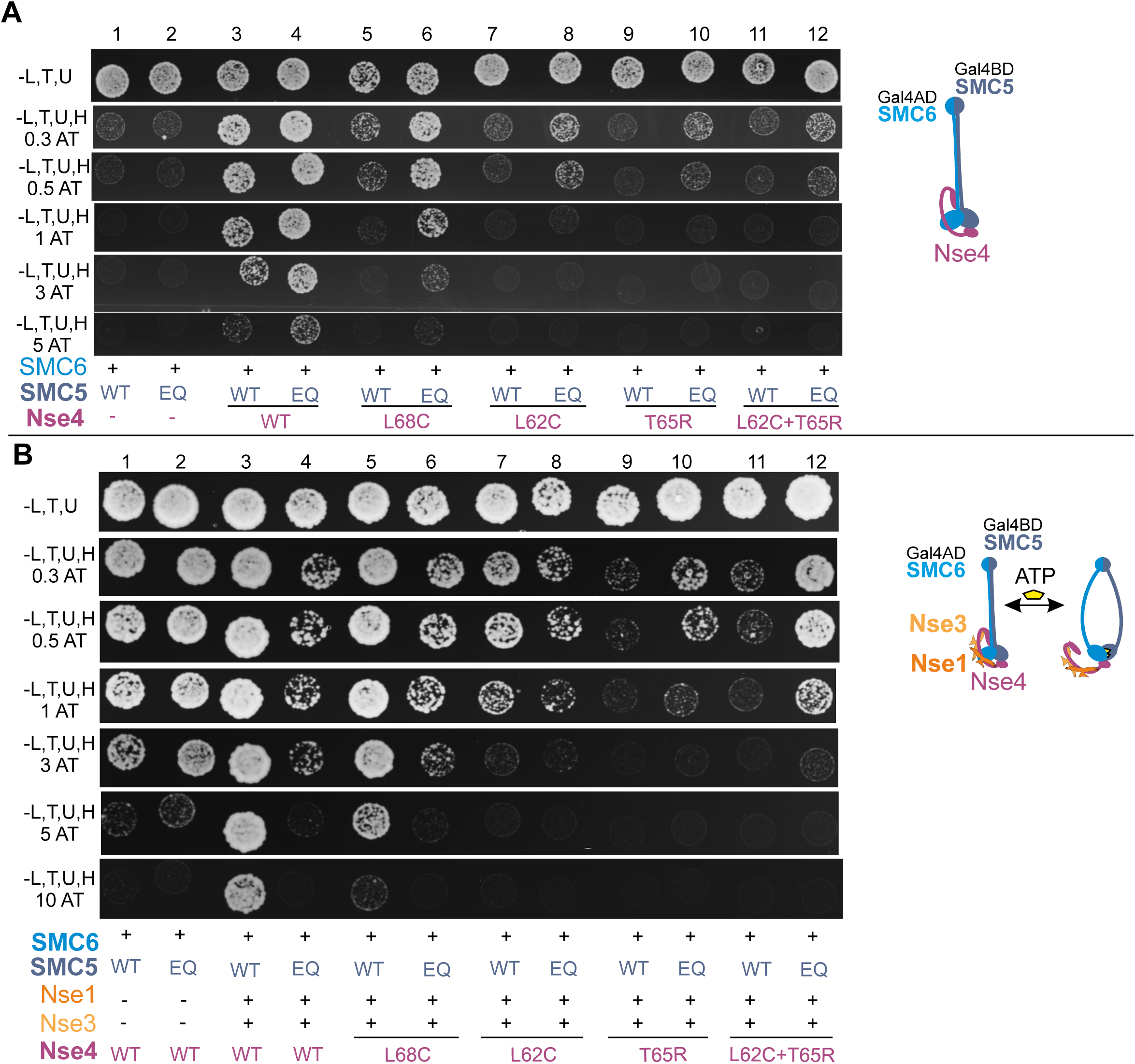
A role of Nse4 and ATP molecules in bridging of SMC5-SMC6. (A) Nse4 is essential for the bridging of the hybrid SMC5-SMC6 constructs (columns 1-4). The SMC5/E995 conserved residue was mutated to glutamine (EQ) to inhibit ATP hydrolysis. The ATP retention has only mild additive effect on the stability of the SMC5-Nse4-SMC6 complex (scored on plates containing increasing concentrations of 3-Amino-1,2,4-triazole). The Nse4 mutations affected the stability of the wild-type SMC5 complexes more dramatically than the stability of the SMC5/E995Q mutant complexes (compare odd and even columns). Wild-type (WT) or E995Q (EQ) mutant versions of SMC5 are labelled in grey below the panels (further details as in Figs. 1 and 2). (B) Addition of the Nse1 and Nse3 KITE proteins to the above SMC5/SMC6/Nse4 system stabilizes the SMC5-Nse4-SMC6 bridge. Although the KITE proteins stabilize the ATP-free SMC5-Nse4-SMC6 complex (columns 1 and 3), they destabilize ATP-bound complex (columns 2 and 4). The Nse4 mutations decrease stability of the ATP-free SMC5-SMC6-Nse4-Nse3-Nse1 complex gradually (columns 5, 7, 9 and 11) while the stability of the ATP-bound complexes drops first (columns 6, 8, and 10) and then it recovers in the L62C, T65R double mutant (column 12).

When we reduced the Nse4 binding affinity to SMC6 using the specific Nse4 mutations described above, the stability of the wild-type SMC5 complexes dropped more dramatically than the stability of the SMC5/E995Q mutant complexes (Fig. 3A, compare odd and even columns). For example, the L68C mutation reduced stability of the wild-type SMC5-Nse4-SMC6 complex significantly while the ATP molecule (in SMC5/E995Q) stabilized the Nse4/L68C complex (Fig. 3A, columns 5 and 6). Further reduction of the Nse4 binding affinity (Fig. 3A, columns 7-12) led to further drops in stability of SMC5-Nse4-SMC6, again, with more stable hydrolytic mutants. Although the stability of the Nse4/L62C, T65R double mutant complex was very low, the residual affinity of Nse4 to SMC6 still supported ATP binding in the SMC5/E995Q mutant (Fig. 3A, columns 11 and 12). These data suggest that ATP contributes significantly to the SMC5-SMC6 bridging when the Nse4 affinity is reduced and that the Nse4 and ATP interactions are synergistic in the SMC5-Nse4-SMC6 complex.

### KITE-bound Nse4 is constrained upon ATP binding

To analyse the role of ATP binding to SMC5-SMC6 within the complex stabilized by KITE proteins, we added Nse1 and Nse3 to the p416ADH1-Nse4 plasmid (p416ADH1-Nse4+Nse3+Nse1 construct, 5Y2H; Fig. 3B; [30, 37]). Consistent with their Nse4-stabilizing roles (Fig. 1B), the KITE proteins increased the stability of the SMC5-Nse4-SMC6 complex significantly (Fig. 3B, compare columns 1 and 3). Surprisingly, addition of the SMC5/E995Q ATP-hydrolysis mutation greatly destabilized the SMC5-SMC6-Nse4-Nse3-Nse1 complex (Fig. 3B, compare columns 3 and 4), suggesting antagonistic roles of ATP and KITE subunits.

To ensure that the instability of the ATP-bound SMC5-SMC6-Nse4-Nse3-Nse1 complex is specifically caused by the ATP-mediated SMC5-SMC6 head bridge, we introduced either SMC6/S1045R mutation disturbing the ATP-mediated SMC5-SMC6 dimerization interface or SMC5/K57I mutation abolishing binding of ATP (Suppl. Fig. S2; [14, 36, 38]). As the SMC6/S1045R mutation suppressed instability caused by the SMC5/E995Q mutation (Suppl. Fig. S2, column 10), we can exclude a direct impact of this mutation on SMC5 (e.g. on SMC5-Nse4 interaction). The observation that both mutations suppressed instability caused by the SMC5/E995Q mutation (Suppl. Fig. S2, column 10 and 12) confirm the notion that ATP-mediated SMC5-SMC6 head dimerization causes SMC5-SMC6-Nse4-Nse3-Nse1 instability.

Given that both ATP and Nse4 bridge SMC5-SMC6 heads, our data suggest that ATP constrains KITE-bound Nse4 bridge (and vice versa; Fig. 3B, compare columns 2 and 4). Therefore, we reduced Nse4 binding affinity to SMC6 (to release the constraint) using the specific Nse4 mutations described above (Figs. 2 and 3A). With decreasing Nse4 affinity, the stability of the ATP-free complexes gradually dropped to its limit (Fig. 3B, columns 5, 7, 9 and 11), suggesting that only Nse4 brought SMC5-SMC6 together. In contrast, the stability of the ATP-bound SMC5-SMC6-Nse4-Nse3-Nse1 complexes dropped first (columns 6, 8, and 10) and then it partially recovered in the L62C, T65R double mutant (column 12). In the single mutants, the reduced Nse4 binding affinity (balance between the Nse4 binding and competing binding of ATP) resulted in decreased stability of these complexes. In the double mutant, residual Nse4 binding was too weak to compete the ATP binding and therefore ATP became the major bridge (manifested as increased stability of the SMC5-SMC6-Nse4-Nse3-Nse1 complex). These data suggest that ATP constrains KITE-bound Nse4 bridge (and vice versa; i.e. the stability of the KITE-containing SMC5/6 complexes depends on the balance between the Nse4 binding and competing binding of ATP).

### The ATP-mediated constraint depends on the KITE subunits

In the SMC5/6 complex, KITE and kleisin subunits form a tight Nse1-Nse3-Nse4 sub-complex mediated by their mutual interactions (Fig. 4A; [27, 35]). As ATP destabilized the Nse4 bridge only in the presence of the KITE subunits (Fig. 3), we introduced mutations specifically affecting stability of the Nse1-Nse3-Nse4 trimer to evaluate a role of the KITE proteins. Specific mutations disturbing only individual Nse1-Nse3 (Nse1/Q18A, M21A) and Nse3-Nse4 (Nse4/del87-91) binary interactions (Fig. 4A, compare columns 3 against 4 and 9 against 10) did not affect the stability of the whole Nse1-Nse3-Nse4 trimer (Fig. 4A, columns 6 and 7), but their combination compromised trimer assembly (Fig. 4A, column 8). When we introduced this combination of Nse1 and Nse4 mutations to the SMC5-SMC6-Nse4-Nse3-Nse1 complex, its stability was reduced as the KITE proteins lost their ability to bind and stabilize Nse4 (Fig. 4B, compare columns 6 and 8). In contrast, when we introduced this combination of Nse1 and Nse4 mutations to the SMC5/E995Q hydrolytic mutant complex, the stability of the ATP-bound complex was increased (Fig. 4B, compare columns 7 and 9), suggesting that the ATP-induced constraint of Nse4 depends on its binding to the KITE subunits. Importantly, there was no difference between the stability of the ATP-free and ATP-bound complexes (compare columns 8 and 9), further corroborating our conclusion that the ATP-mediated constraint depends on the binding of KITE dimer to Nse4. Furthermore, the Nse4/del87-91 mutation compromising only Nse3-Nse4 interaction (Figs. 4A, columns 4 and 6) had a suppressing effect on SMC5/E995Q complex similar to the double mutant (Fig. 4B, compare columns 9 and 11), suggesting that the binding of Nse3 to the Nse4 linker partially constrained it. Our data suggest that the instability of the SMC5/6 complex induced by ATP binding is dependent on the binding of KITE proteins to the Nse4 kleisin linker.

**Figure 4.**
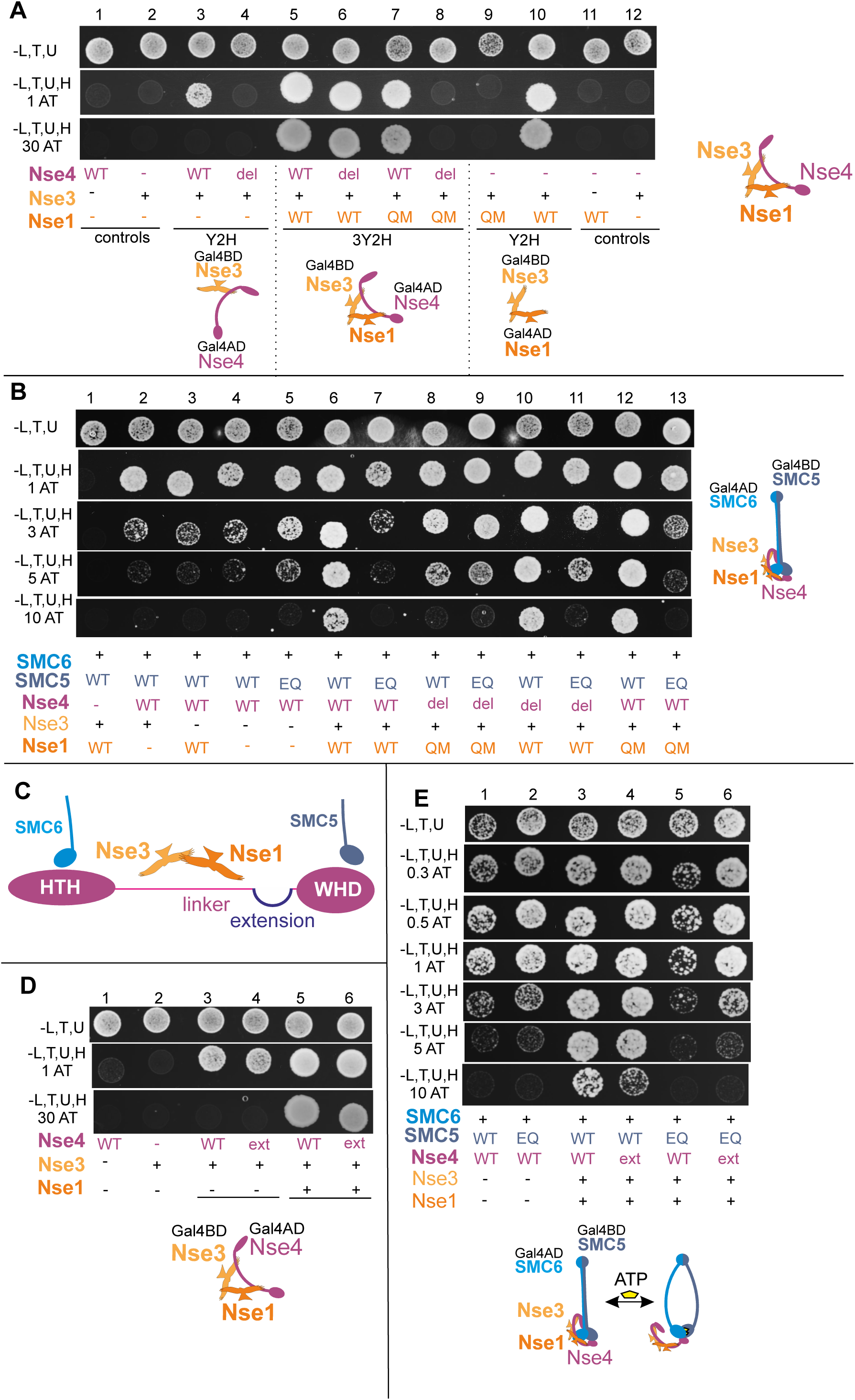
KITE proteins constrain Nse4 kleisin linker. (A) The Nse1-Nse3-Nse4 subcomplex is held by mutual interactions between its subunits. The Nse4/del87-91 (del) deletion disturbs the Nse3-Nse4 binary interaction (columns 3 and 4) and the Nse1/Q18A, M21A (QM) double mutation abrogates the Nse1-Nse3 binary interaction (columns 9 and 10), but they do not alter stability of the Nse1-Nse3-Nse4 trimer individually (columns 6 and 7). However, combination of these two mutations reduces the stability of Nse1-Nse3-Nse4 significantly (column 8). (B) The combination of the Nse4/del87-91 and Nse1/Q18A, M21A mutations compromises stability of the wild-type SMC5-SMC6-Nse4-Nse3-Nse1 complex (compare columns 6 and 8), but increases the stability of the ATP-bound complex (compare columns 7 and 9). Notably, there is no difference between stability of ATP-free and ATP-bound complexes (compare columns 8 and 9), suggesting that the binding of KITE proteins to Nse4 destabilizes ATP-bound complexes. Furthermore, the Nse4/del87-91 mutation alone also partially supresses the instability of the ATP-bound complex (column 11), suggesting that the binding of Nse3 to Nse4 linker constrains the linker. (C) Schematic of the Nse4 regions with their binding partners (depicted above). The 30 amino acid extension is inserted at the end of the linker. (D) The 30 amino acid extension (ext) was tested in control experiments for the stability of either Nse3-Nse4 binary interaction (column 4) or Nse1-Nse3-Nse4 trimer (column 6). (E) The extension of the linker results in partial relieve of the ATP-induced tension in the SMC5/E995Q complex (compare columns 5 and 6).

### The limited size of the Nse4 linker poses mechanical constraint

The KITE dimers bind linker regions of kleisin molecules in SMC complexes [7, 16, 28, 39–41]. To explore the role of the Nse4 linker in propagation of the ATP-induced constraint, we inserted a 30 amino acid extension at the putative end of the linker (Fig. 4C; [27]) to lengthen the linker. The Nse4 extended construct bound Nse1-Nse3 KITE proteins normally (Fig. 4D) and formed the SMC5-SMC6-Nse4-Nse3-Nse1 complex with the stability similar to that with the normal Nse4 construct (Fig. 4E, columns 3 and 4). Interestingly, combination of the Nse4 extended construct with the SMC5/E995Q ATP-hydrolysis mutant increased the stability of the SMC5-SMC6-Nse4-Nse3-Nse1 complex, suggesting that the extended Nse4 linker partially alleviated ATP-induced constraint (Fig. 4E, columns 5 and 6).

Taken together, we propose a model in which the KITE proteins shape the kleisin linker connecting SMC heads. The KITE-shaped Nse4 linker fits the ATP-free conformation of SMC5/6 (and therefore increases its stability), while the ATP-bound conformation is not compatible with the KITE-shaped Nse4 linker (and therefore constrains the Nse4 bridge; Fig. 5). This model suggests a key role of the kleisin and KITE subunits in molecular mechanisms driving the SMC5/6 dynamics.

**Figure 5.**
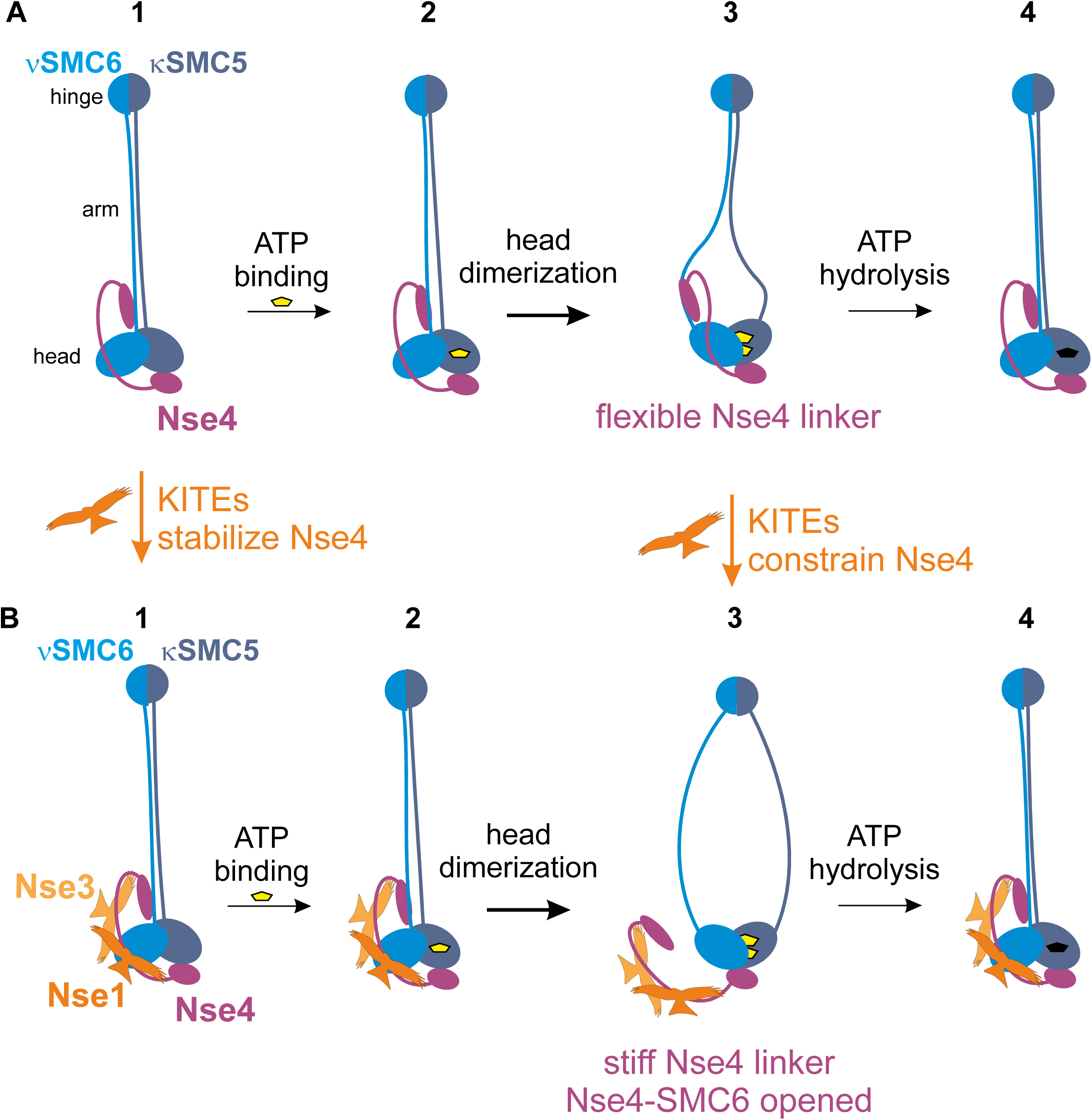
SMC5/6 ATPase cycle model. Binding of ATP to the SMC heads leads to their dimerization and changes their mutual position (states 1-3). (A) Nse4 bridges SMC heads and supports the ATP-mediated head dimerization in the absence of the KITE proteins. (B) Addition of the Nse1 and Nse3 KITE proteins stabilizes the otherwise flexible Nse4 linker in a position that is favoured in the ATP-free complex (state B1). In contrast, the KITE-bound Nse4 linker shape is adverse in the ATP-bound complex (state B3). The stiff KITE-bound Nse4 linker transduces a pulling force generated by the binding of ATP to the SMC5-SMC6 heads and forces the Nse4-SMC6 interface apart (state B3). After ATP hydrolysis, the SMC5-SMC6 head dimer is dissolved and Nse4 is reattached to the SMC6 arm (state B4).

## DISCUSSION

The kleisin subunits bridge the SMC heads in an asymmetric way and lock the SMC ring at its head side [12, 13]. We have shown that Nse4 belongs to the kleisin superfamily of proteins and binds strongly to κSMC5 head via its Nse4 C-terminal WHD [27]. However, we observed only weak binding of Nse4 to νSMC6 [27] and other studies actually failed to show the Nse4-SMC6 interaction [31, 32, 42]. Here we developed several systems to prove and analyse the interaction between Nse4 and SMC6. We mapped the Nse4-SMC6 interface in detail and found that the Nse4-SMC6 interaction mode is similar to the other νSMC-kleisin interactions [16–18, 26, 34]. Therefore, we assume that Nse4 bridges SMC5-SMC6 proteins in a way similar to kleisins in the other SMC complexes, except that the Nse4 bridge is specifically modulated by the Nse1-Nse3 KITE subunits in the SMC5/6 complex (see below).

The kleisins lock the SMC rings that can embrace DNA or extrude a loop in an ATP-dependent way [43]. To release such entrapped DNA (or extrude the loop), the SMC-SMC or SMC-kleisin interface must be open. It was proposed that the νSMC-kleisin interface opens and serves as an exit gate for trapped DNA [19, 44, 45]. In the cohesin complex, the Scc1-SMC3 interface is opened upon ATP binding (in the presence of the Pds5 and Wapl regulators; [19–24]). Our data show that the SMC5/6 complex is unstable in the ATP-bound state, suggesting that one or more interfaces are compromised upon ATP binding (Fig. 3B). Given the weak nature of the Nse4-SMC6 interaction, we assume that this interaction is prone to dissociation and that the Nse4-SMC6 interface is opened upon ATP binding. Consistent with the latter notion, the ATP-mediated constraint was released when the Nse4-SMC6 interaction was disturbed (Fig. 3B). Therefore, we hypothesize that the binding of ATP to the SMC5-SMC6 heads constrains the Nse4 bridge (when bound by the Nse1 and Nse3 KITE subunits; see below and Fig. 5) and this constraint is released via dissociation of the Nse4-SMC6 interaction (Fig. 3B).

The ATP binding induces changes in the mutual positions (and conformations) of the SMC heads and arms [10, 26, 46–48]. Consequently, the shape of the whole SMC complex changes from a rod-like conformation (with juxtaposed arms stabilized by their mutual interactions) to the less stable open ring (Fig. 5). At least part of the ATP-induced SMC5/6 instability (Fig. 3B) might be a consequence of the shape transition from rod to ring (our electron microscopy data suggest that the human SMC5/6 complex can adopt both shapes; M. Adamus, unpublished data). However, we observed this instability only in the presence of the KITE subunits, suggesting that either our system is unable to detect the SMC5-SMC6 rod-to-ring shape transition (and monitors only Nse4 bridge opening) or the KITE proteins are required for the full rod-to-ring shape transition. The latter possibility is consistent with the prokaryotic SMC/ScpAB data which suggest an important function of the KITE subunits in pulling SMC arms [25, 26]. In our model, the binding of ATP to the SMC5-SMC6 heads pulls their arms apart and constrains the KITE-bound Nse4 bridge (Fig. 5).

It was shown that kleisin-interacting proteins (particularly KITE and HAWK subunits) bind and shape linker regions of the kleisin molecules [7, 11, 16, 28, 39–41, 49–51]. Our data showed that the binding of the Nse1-Nse3 KITE dimer increases the Nse4 ability to bind SMC5-SMC6 subunits in ATP-free state (Figs. 1B and 3B), suggesting that the KITE binding may shape Nse4 to fit the ATP-free conformation of the SMC5/6 complex. In contrast, the KITE-bound Nse4 linker is barely compatible with the ATP-bound conformation of SMC5/6. Consistent with these notions, 30 amino acid long extension of the Nse4 linker partially relaxed its stiff shape and resulted in a mild drop in the stability of the ATP-free complex (Fig. 4E). In contrary, this extension partially released the ATP-induced constraint in the ATP-bound complex. Further extension of the Nse4 linker decreased the stability of the ATP-free complex further (as the linker became more flexible) to the levels similar to the ATP-bound complex (90 amino acid extension fully released ATP-induced constraint; L. Vondrova, unpublished data). Therefore, we suggest that KITE subunits shape the Nse4 linker to fit the ATP-free complex optimally and to facilitate opening of the complex upon ATP binding (Fig. 5). Consistent with this conclusion, the ATP-mediated constraint was partially suppressed upon release of the part of the Nse4 linker from the Nse3 WHB pocket (Fig. 4B; [28, 35]). Altogether, we hypothesize that the Nse4 linker is stiffened upon its KITE binding and transduces a pulling force generated by the binding of ATP to the SMC5-SMC6 heads (Fig. 5B). In consequence, the Nse4-SMC6 interface opens and releases the Nse4 constraint. After ATP hydrolysis, the SMC5-ATP-SMC6 head dimer is dissolved and Nse4 is reattached to the SMC6 neck.

Similarly, it was proposed that binding of the HAWK (Scc3 and Pds5) and Wapl proteins to Scc1 kleisin stiffens its linker region and transduces conformational energy of the ATP-dependent SMC head dimerization to the dissociation of Scc1 from Smc3 [19, 45]. In the absence of the Pds5-Wapl regulators, the cohesin’s head movements driven by ATP binding and hydrolysis cannot be effectively coupled to exit gate opening as the Scc1 linker is flexible. Interestingly, the size of the Scc1 linker is much longer (cca 400 amino acids) than the size of the Nse4 linker (and ScpA linker; both having cca 100 amino acids; J. Palecek, unpublished data). Accordingly, the Scc3 HAWK subunit covers only a small part of the Scc1 linker and requires Pds5-Wapl regulators to shape the long Scc1 linker while the KITE subunits are sufficient to shape their short kleisin partners. As mentioned above, the extension of the Nse4 linker region suppressed the ATP-induced constraint, suggesting that the short size of the Nse4 linker is critical for the dynamics of the SMC5/6 complex. Consistent with this notion, the integration of the 30 amino acid long extension to the genomic copy of the fission yeast Nse4 resulted in severe DNA repair phenotypes (L. Vondrova, unpublished data).

As the KITE subunits are stable components of the SMC complexes, we assume that the openings of the kleisin-νSMC interfaces are intrinsically coupled to their ATP cycles. It was proposed for the SMC/ScpAB complex that the opening of the kleisin-νSMC interface might be a part of its ATPase cycle generating loops along DNA strands [52, 53]. The SMC5/6 data are also consistent with the above notions as the ATPase activity of the SMC5/6 complex is needed for its topological binding [36]. Interestingly, the Nse1-Nse3 KITE subunits bind DNA and this interaction could anchor or transduce DNA during the loop extrusion mediated by the SMC5/6 complex [37]. In contrast, dissociation of the kleisin-νSMC interface is tightly controlled by Pds5-Wapl regulators and is coupled to cohesin release from chromosome arms. Consistent with these differences, the Scc1-SMC3 fusion is tolerated in cells (as they can release cohesin by other ways; [20, 21, 23, 54]) while the fusion of Nse4 and SMC6 is lethal in fission yeast (L. Vondrova, unpublished data). Our results suggest very similar mechanics shared by the prokaryotic SMC/ScpAB and eukaryotic SMC5/6 complexes (while distinguishing them from cohesin) and further support our recently proposed close evolutionary relationship between these complexes [7].

However, there are also apparent differences between the SMC/ScpAB and SMC5/6 complexes. Particularly, the Nse1 KITE subunit contains an RING-finger ubiquitin ligase domain which may add a specific regulatory level to SMC5/6 [55]. Interestingly, our *in vitro* and *in vivo* experiments showed an Nse1-dependent Nse4 kleisin ubiquitination of its linker both in *S. pombe* and human proteins ([56]; P. Kolesar, unpublished data). Such a bulky post-translational modification on the Nse4 linker could alter its binding to KITE partners or its stiffness (i.e. alter the SMC5/6 ring opening). Altogether, similarities and differencies between SMC complexes at different levels of their architecture may stay behind their different functions in genomes and remain an intriguing avenue of future research.

## MATERIAL AND METHODS

### Plasmids

Most of the Y2H constructs were prepared previously: pGBKT7-Nse3(aa1-328), pGADT7-Nse3(aa1-328) and pGBKT7-Nse4(aa1-300) constructs were created in [33], pOAD-Nse1(aa1-232) was created in [35], pGADT7-SMC6 (aa1-1140) was described in [37]. To generate pGBKT7-SMC5(aa1-1076) construct, the yeast *S.pombe* SMC5 cDNA was PCR amplified by oLV511+oLV486 (Supplementary Table 2) and inserted into the *NcoI–SalI* digested pGBKT7 by In-Fusion cloning protocol (Clontech). Nse5 was cloned into pGBKT7 vector using *NcoI* and *Sa*lI sites and classical T4 ligase protocol.

To create the pGADT7-Nse4(aa1-300)/WT, pGADT7-Nse4(aa1-300)/del87-91 and pGADT7-Nse4(aa1-300)/ext constructs, Nse4 was PCR amplified from the corresponding p416ADH1-Nse4 plasmids (see below) by oLV575+oLV576 and inserted into *NdeI/BamHI* digested pGADT7 by In-Fusion cloning protocol.

Multicomponent Y2H system was described in the protocol book series Methods in Molecular Biology [30]. The p416ADH1-Nse1(aa1–232) construct was created previously [35]. Construction of p416ADH1-Nse4 (aa1-300), p416ADH1-Nse3(aa1–328)+Nse4(aa1–300) and p416ADH1-Nse3(aa1–328)+Nse4(aa1–300)+Nse1(aa1–232) was described previously ([30, 37]; the vector name pPM587 equals to p416ADH1). p416ADH1-Nse4(aa1–300)+Nse1(aa1–232) and p416ADH1-Nse3(aa1–328)+Nse1(aa1–232) were prepared by PCR-amplification of ADH1-Nse1(1-232)-CYC terminator from p416ADH1-Nse1 by KB353+KB354 and its insertion into *KpnI* digested p416ADH1-Nse4 (aa1-300) and p416ADH1-Nse3(aa1–328) by In-Fusion protocol, respectively. The p416ADH1-Nse6(1-522) construct was prepared using PCR-amplification of Nse6 by EB77 + EB78 primers and insertion into *SalI/SpeI* digested p416ADH1 by In-Fusion protocol.

To prepare p416ADH1-Nse4-ext, *SalI* restriction site in the p416ADH1 multi-cloning site (MCS) was mutated by site-directed mutagenesis (SDM; see below) with oLV522 + oLV523. Then, *SalI* site was inserted by SDM behind the aa174 with oLV520 + oLV521. This construct was *SalI* digested, (G_4_S)_6_ linker was amplified by oLV579 + oLV580 and inserted using the In-Fusion cloning protocol.

Construct for the *S. pombe* genome integration was prepared as follows: 1. Nse4 was cloned within the *BamHI* and *EcoRI* sites of the pSK-ura4 plasmid; 2. genomic sequence downstream of the Nse4 gene was PCR amplified (JP414 and JP415) and inserted to the pGEM-Easy vector (Promega); 3. The *SacI-SalI* fragment of the pSK-Nse4-ura4 plasmid was inserted to the *SacI*-*XhoI* digested pGEM-3’end construct to get the pGEM-Nse4(WT)-ura4 integration plasmid. To create pGEM-Nse4(T65R)-ura4 and pGEM-Nse4(L62C,T65R)-ura4, the Nse4 sequences were PCR amplified from p416ADH1-Nse4 mutant constructs by oLV680 + oLV681 (Supplementary Table 2) and inserted into *BamHI-EcoRI* digested pGEM-Nse4(WT)-ura4 integration construct by the In-Fusion cloning protocol.

### Site-directed mutagenesis

The QuikChange Lightning Site-Directed Mutagenesis Kit (Agilent Technologies) was used to create mutations in pGBKT7-SMC5, pGADT7-SMC6, p416ADH1-Nse4, p416ADH1-Nse3+Nse4+Nse1, pGADT7-Nse4, pOAD-Nse1, p416ADH1-Nse1, and pGEM-Nse4. The sequences of primers used for mutagenesis are listed in Supplementary Table 3.

### Yeast two-hybrid assays

The Gal4-based Y2H system was used to analyze *S. pombe* SMC5/6 complex interactions (detailed protocols are described in the book series Methods in Molecular Biology [30]). Briefly, three plasmids pGBKT7, pGADT7 and p416ADH1 with corresponding proteins were co-transformed into the *Saccharomyces cerevisiae* PJ69–4a strain and selected on SD -Leu, - Trp, -Ura plates. Drop tests were carried out on SD -Leu, -Trp, -Ura, -His (with 0; 0.3; 0.5; 1; 2; 3; 4; 5; 10; 15; 20; 30 mM 3-aminotriazole) plates at 28°C. Each combination was co-transformed at least three times and at least three independent drop tests were carried out.

### Generation of *nse4* mutant strains of *S. pombe*

Standard genetic techniques were used for preparation of fission yeast *S. pombe* diploid strain by crossing *ade6-M216* strain with *ade6-M210* [57]. The pGEM-Nse4-ura4 wild-type or mutant integration constructs (digested by *BamHI* and *SalI*) were transformed into the diploid strain and the transformants were selected on –ura –ade plates. Using standard genetic techniques, spores of the *nse4+/nse4-mutant* strains were generated and dissected.

## PEPSCAN-ELISA

Was performed as described previously [28] with a peptide library (Mimotopes, Australia) of the aa 875-1024 region of the *S. pombe* Smc6 protein (Supplementary Table 1). The library was linked to biotin via an additional peptide spacer of serine–glycine–serine–glycine. The peptides were prebound to ELISA plates (coated with streptavidin) and washed three times with binding buffer (PBS with 0.5% Nonidet NP40). Then, *S. pombe* His-S-Nse4 (aa1-150) protein was added and incubated overnight. Unbound protein was washed (three times) and the peptide-bound Nse4 protein was quantified using anti-His (Sigma H1029, 1:10000) and anti-mouse HRP-conjugated (Sigma A0168, 1:10000) antibodies, respectively. *H. sapiens* His-TRF2 (aa1-542) protein was used as a negative control in the same way [29].

## SUPPORTING INFORMATION

**Supplementary Figure S1. Nse4 binds neck region of SMC6.**

Peptide library (A) and multi-component yeast two-hybrid systems (C-E) were employed to determine the SMC6 region and residues binding to Nse4. (A) Quantification of relative binding of the Nse4(1-150) protein (Nse4; red columns) to the SMC6 synthetic peptides (listed in Supplementary Table 1) using the PEPSCAN-ELISA method. The SMC6(aa960-984) peptide exhibits the highest affinity and specificity to Nse4. Results show mean ± SEM of 3 independent measurements. His-TRF2 protein (TRF2; white column) was used in the control experiment. (B) Alignment of the C-terminal SMC6 neck region. The orthologs are from *Schizosaccharomyces pombe* (*S.p.*), *Aspergillus nidulans* (*A.n.*), *Aspergillus clavatus* (*A.c.*), *S. cerevisiae* (*S.c.*), *Danio rerio* (*D.r.*), *Xenopus laevis* (*X.l.*), *Ornithorhynchus anatinus* (*O.a.*), *Loxodonta africana* (*L.a.*), *Monodelphis domestica* (*M.d.*), *Dasypus novemcinctus* (*D.n.*), *Mus musculus* (*M.m.*), *Homo sapiens* (*H.s.*). “+”, mutation not affecting SMC6 interactions; “-“, mutation disrupting all SMC6 complexes; red minus, mutation specifically disrupting the Nse4-SMC6 interaction. Amino acid shading represents following conserved amino acids: *dark green*, hydrophobic and aromatic; *light green*, polar; *pink*, basic. (C-E) To identify Nse4-binding residues, stability of the following SMC6 mutant complexes was tested: SMC6-Nse4-Nse3-Nse1 (C), SMC6-Nse4-SMC5 (D) and SMC6-Nse5-Nse6 (E). (C) The full-length hybrid SMC6 (fused to Gal4AD domain) and Nse4 (fused to Gal4BD domain) were co-transformed together with Nse1-Nse3 construct (p416ADH1 vector) into PJ69 cells. Formation and stability of the SMC6-Nse4-Nse3-Nse1 complex was scored by growth of yeast PJ69 transformants on plates without Leu, Trp, Ura and His, containing 0.3 mM 3-Amino-1,2,4-triazole (-L,T,U,H, 0.3AT panel). (D) Similarly, the full-length SMC6 (fused to Gal4AD domain) and SMC5 (fused to Gal4BD domain) were co-transformed together with Nse4 full-length construct (in p416ADH1 vector) and stability of the SMC5-Nse4-SMC6 complex was scored on plates containing 0.5 mM 3-Amino-1,2,4-triazole (-L,T,U,H, 0.5AT panel). The L964A, L965A, L968A, E969A, L972A and R975A mutations reduce stability of the SMC6-Nse4 complexes (C and D). (E) In the control experiment, the same mutations were introduced to SMC6-Nse5-Nse6 complex (constituted of the full-length Gal4AD-SMC6, Gal4BD-Nse5 and non-hybrid Nse6). Stability of the SMC6-Nse5-Nse6 complex was scored on plates containing 3 mM 3-Amino-1,2,4-triazole (-L,T,U,H, 3AT panel). The L964A, L968A and E969A mutations affected all SMC6 complexes (C-E). In contrast, the highly conserved L965, L972 and L975 SMC6 residues (panel B) were required specifically for binding to Nse4. Wild-type (WT) or mutant versions of SMC6 are labelled in blue below the panels; “-“, control empty vector; “+”, co-transformed construct (as labelled at the left side). Growth of the transformants was verified on control plates without leucine, tryptophan and uracil (-L,T,U). All Y2H tests were repeated at least 3 times.

**Supplementary Figure S2. The SMC5-SMC6-Nse4-Nse3-Nse1 instability is dependent on the ATP binding and SMC5-SMC6 head dimerization**.

(A-B) Alignment of the conserved SMC head motifs mutated in the SMC5 and SMC6 constructs. Arrows point to the positions of the fission yeast SMC5/K57I mutation in the Walker A motif disturbing ATP binding (A), the SMC6/S1045R mutation in the Signature motif abrogating ATP-mediated SMC5-SMC6 head dimerization (B), and the SMC5/E995Q mutation in the Walker B motif inhibiting ATP hydrolysis (B). The SMC homologs are from *Bacilus subtilis* (*B.s.*), *Schizosaccharomyces pombe* (*S.p.*), *Homo sapiens* (*H.s.*). Amino acid shading as in Fig. S1. (C) Stability of the SMC5-SMC6-Nse4 (columns 1-6) and SMC5-SMC6-Nse4-Nse3-Nse1 (columns 7-12) complexes was scored on plates containing increasing concentrations of 3-Amino-1,2,4-triazole (AT). The SMC5/K57I (KI) mutation disturbing ATP binding and SMC6/S1045R (SR) mutation abrogating ATP-mediated SMC5-SMC6 head dimerization have no effect on the SMC5-SMC6-Nse4 complex irrespective of the SMC5/E995Q mutation (columns 3-6). In contrast, destabilizing effect of the SMC5/E995Q mutation was fully supressed by both KI and SR mutations in the SMC5-SMC6-Nse4-Nse3-Nse1 complex (compare columns 8, 10 and 12), suggesting that the instability is caused by the ATP binding and ATP-mediated head dimerization. Mutant versions of SMC5 and SMC6 are labelled in grey and blue, respectively (further details as in Fig. S1).

Supplementary Table 1: SMC6(aa875-1024) peptide library

Supplementary Table 2: Primers used for PCR

Supplementary Table 3: Primers used for site-directed mutagenesis

## ACKNOWLEDGMENTS

We thank C. Hofr for providing the purified His-hTRF2 protein for the ELISA assays. We are grateful to A.R. Lehmann and K. Zabrady for the critical reading of our manuscript.

