## Supplementary material for "A role of the Nse4 kleisin and Nse1/Nse3 KITE subunits in the ATPase cycle of SMC5/6": Supplemntary information

Suppl. Figure 1

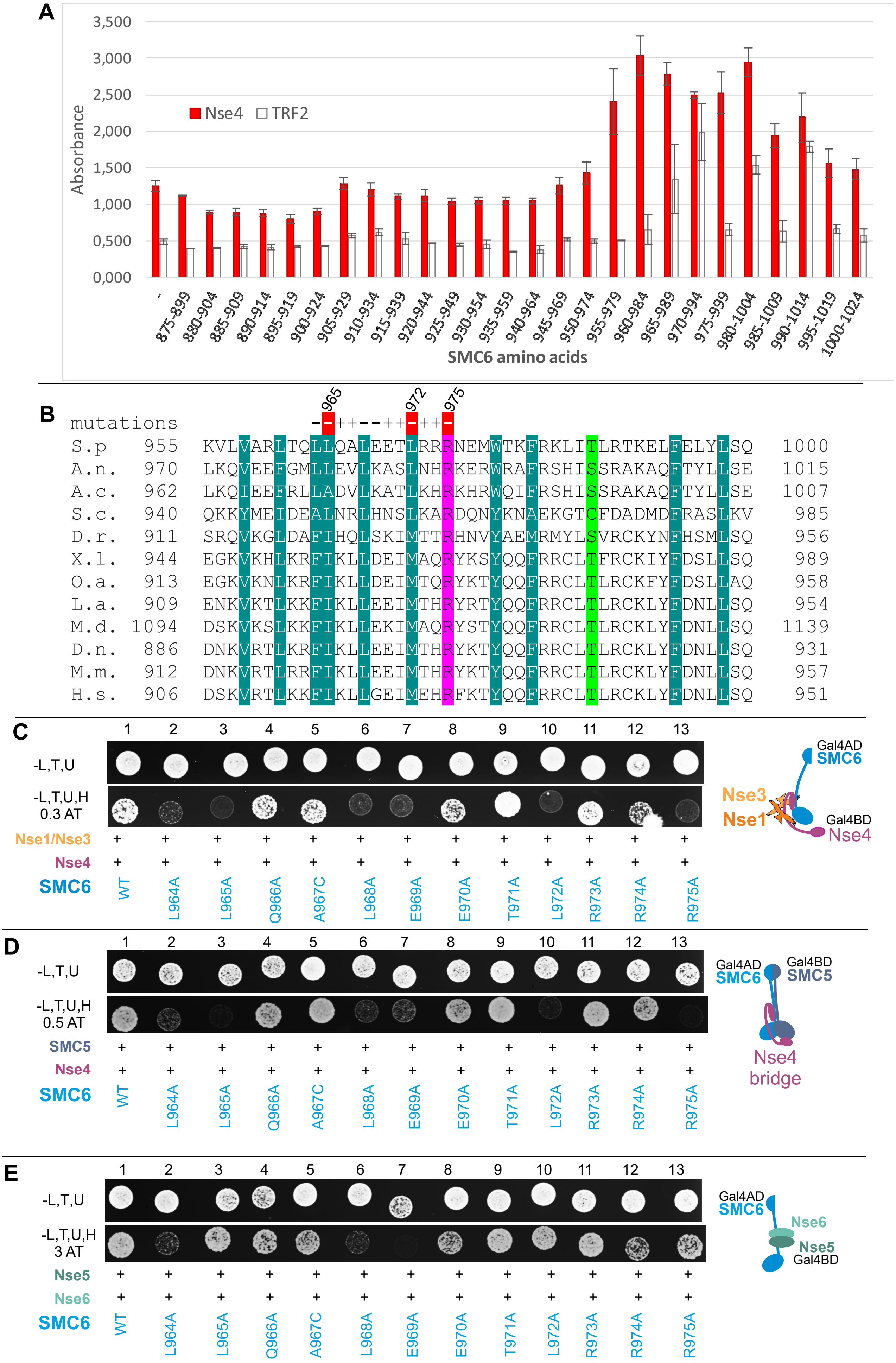

Suppl Figure 2

**A**

```
                          spSMC5/K57I
                               ↓
Smc   B.s.   27   TAVVGPNGSGKSNITDAI   45
Smc1  S.p.   28   TSIIGPNGAGKSNLMDAI   46
Smc2  S.p.   28   NAITGLNGSGKSNILDAI   46
Smc3  S.p.   28   NVIVGRNGSGKSNFFAAI   46
Smc4  S.p.  151   SSIVGPNGSGKSNVIDAL  169
Smc5  S.p.   47   NLIIGPNGTGKSTIVSAI   65
Smc6  S.p.  120   NFVIGHNGSGKSAILTGL  138
Smc1  H.s.   28   TAIIGPNGSGKSNLMDAI   46
Smc2  H.s.   28   NAITGLNGSGKSNILDSI   46
Smc3  H.s.   28   NVIVGRNGSGKSNFFYAI   46
Smc4  H.s.  109   SCIIGPNGSGKSNVIDSM  127
Smc5  H.s.   76   NMIVGANGTGKSSIVCAI   94
Smc6  H.s.   72   NFVVGNNGSGKSAVLTAL   90
                    : * **.***  .   :
```

**B**

```
                         spSMC6/S1045R                               spSMC5/E995Q
                              ↓                                            ↓
Smc   B.s.  1087   LLSGGERALTAIALLFSILKVRPVPFCVLDEVEAALDE   1125
Smc1  S.p.  1130   QLSGGEKTMAALALLFAIHSYQPSPFFVLDEIDAALDQ   1168
Smc2  S.p.  1084   ELSGGQRSLVALALIMSLLKYKPAPMYILDEIDAALDL   1122
Smc3  S.p.  1095   QLSGGQKSLCALTLIFAIQRCDPAPFNILDECDANLDA   1133
Smc4  S.p.  1228   NLSGGEKTLSSLALVFALHNYKPTPLYVMDEIDAALDF   1266
Smc5  S.p.   964   RQSGGERSVSTIMYLLSLQGLAIAPFRIVDEINQGMDP   1002
Smc6  S.p.  1042   GLSGGEKSFATICMLLSIWEAMSCPLRCLDEFDVFMDA   1080
Smc1  H.s.  1126   NLSGGEKTVAALALLFAIHSYKPAPFFVLDEIDAALDN   1164
Smc2  H.s.  1083   ELSGGQRSLVALSLILSMLLFKPAPIYILDEVDAALDL   1121
Smc3  H.s.  1113   QLSGGQKSLVALALIFAIQKCDPAPFYLFDEIDQALDA   1151
Smc4  H.s.  1189   NLSGGEKTLSSLALVFALHHYKPTPLYFMDEIDAALDF   1227
Smc5  H.s.   989   HQSGGERSVSTMLYLMALQELNRCPFRVVDEINQGMDP   1027
Smc6  H.s.   985   ALSGGERSFSTVCFILSLWSIAESPFRCLDEFDVYMDM   1023
                     ***::..  :  :: :        *:  .**     :*
```

**C**

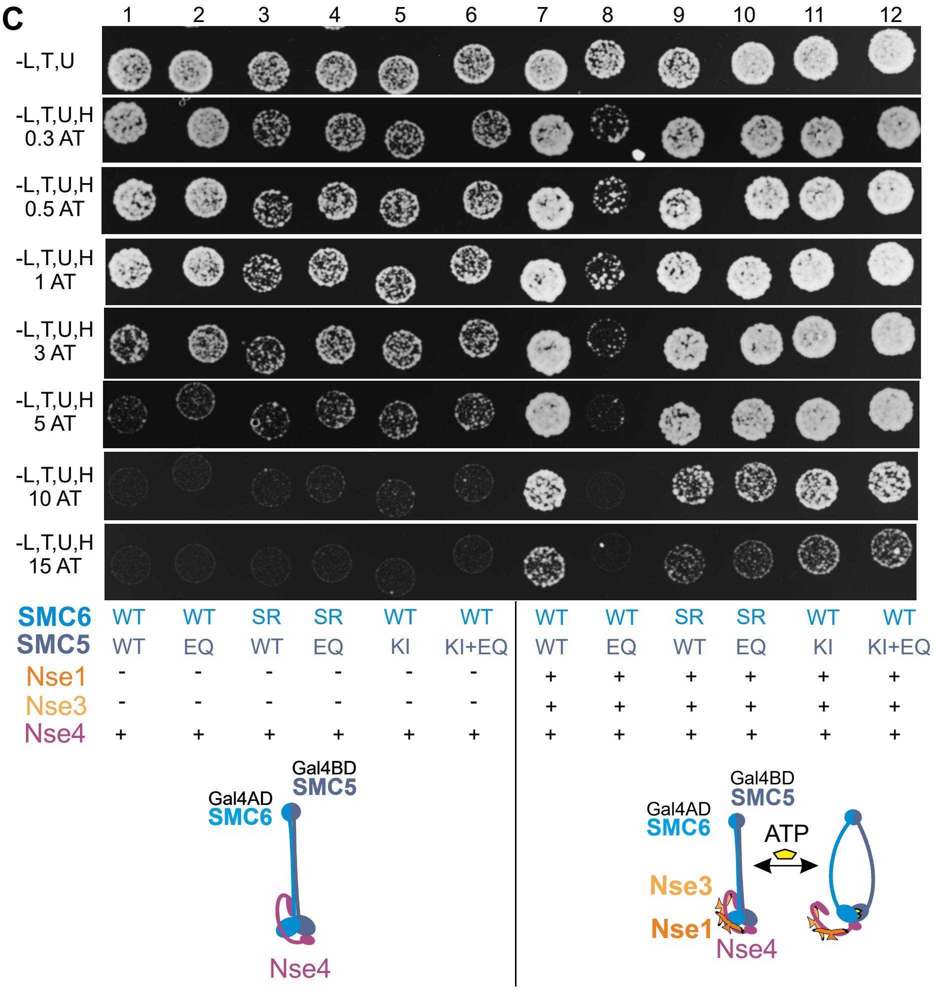

### Supplementary Table 1: SMC6(aa875-1024) peptide library

| aa | Smc6 peptide sequence |
|---|---|
| 875-899 | TNILREKEAKKVQCAQVVADYTAKA |
| 880-904 | EKEAKKVQCAQVVADYTAKANTRCE |
| 885-909 | KVQCAQVVADYTAKANTRCERVPVQ |
| 890-914 | QVVADYTAKANTRCERVPVQLSPAE |
| 895-919 | YTAKANTRCERVPVQLSPAELDNEI |
| 900-924 | NTRCERVPVQLSPAELDNEIERLQM |
| 905-929 | RVPVQLSPAELDNEIERLQMQIAEW |
| 910-934 | LSPAELDNEIERLQMQIAEWRNRTG |
| 915-939 | LDNEIERLQMQIAEWRNRTGVSVEQ |
| 920-944 | ERLQMQIAEWRNRTGVSVEQAAEDY |
| 925-949 | QIAEWRNRTGVSVEQAAEDYLNAKE |
| 930-954 | RNRTGVSVEQAAEDYLNAKEKHDQA |
| 935-959 | VSVEQAAEDYLNAKEKHDQAKVLVA |
| 940-964 | AAEDYLNAKEKHDQAKVLVARLTQL |
| 945-969 | LNAKEKHDQAKVLVARLTQLLQALE |
| 950-974 | KHDQAKVLVARLTQLLQALEETLRR |
| 955-979 | KVLVARLTQLLQALEETLRRRNEMW |
| 960-984 | RLTQLLQALEETLRRRNEMWTKFRK |
| 965-989 | LQALEETLRRRNEMWTKFRKLITLR |
| 970-994 | ETLRRRNEMWTKFRKLITLRTKELF |
| 975-999 | RNEMWTKFRKLITLRTKELFELYLS |
| 980-1004 | TKFRKLITLRTKELFELYLSQRNFT |
| 985-1009 | LITLRTKELFELYLSQRNFTGKLVI |
| 990-1014 | TKELFELYLSQRNFTGKLVIKHQEE |
| 995-1019 | ELYLSQRNFTGKLVIKHQEEFLEPR |
| 1000-1024 | QRNFTGKLVIKHQEEFLEPRVYPAN |

**Supplementary Table 2: Primers used for PCR**

| | | | |
|---|---|---|---|
| SMC5 In-Fusion pGBKT7 | oLV511 | fw | CTGCATATGGCCATGGATGGCCTTAGGCCT |
| | oLV486 | rev | GCCGCTGCAGGTCGACTTATGACGAAGAAATGAGTGC |
| Nse6 In-Fusion p416ADH1 | EB77 | fw | CCGCTCTAGAACTAGTATGAATGCGTCTAATAACATTTCAA |
| | EB78 | rev | ATGACTCGAGGTCGACTTATCTTTTGTACGCTTGCC |
| Nse4 In-Fusion pGADT7 | oLV575 | fw | CAGATTACGCTCATATGTCCTCCATTGATAAACG |
| | oLV576 | rev | CGAGCTCGATGGATCCTCAGCCATACCAAGTATTACTGT |
| insertion into p416ADH1 | KB353 | fw | CTTTAATTTGCGGCCGGGAGCTCGCCGGGATC |
| | KB354 | rev | CTATAGGGCGAATTGG |
| $(G_4S)_6$ linker insertion | oLV579 | fw | CATTACTACCGTCGAaGGTGGAGGAGGCTCT |
| | oLV580 | rev | TATTTTCTTGGTCGACTGACCCTCCGCCT |
| 3'end of Nse4 | JP414 | fw | CTCGAGATAACGCTTCGTAATTAAGAT |
| | JP415 | rev | GTCGACCTTGCATAAATACTTAGTCC |
| Nse4 In-Fusion pGEM | oLV680 | fw | TAGAACTAGTGGATCCATGTCCTCCATTGATAAACG |
| | oLV681 | rev | GCTTGATATCGAATTCTCAGCCATACCAAGTATTACT |

### Supplementary Table 3: Primers used for site-directed mutagenesis

| | | | |
|---|---|---|---|
| SalI mutation in MCS | oLV522 | fw | TCAAGCTTATCGATACCGTgGACCTCGAGTCATGTAATTAGT |
| | oLV523 | rev | ACTAATTACATGACTCGAGGTCcACGGTATCGATAAGCTTGA |
| SalI insertion into Nse4 | oLV520 | fw | CTGAATGAACGTAACATTACTACCgtcgacCAAGAAAATAACACCACTAAAAATG |
| | oLV521 | rev | CATTTTTAGTGGTGTTATTTTCTTGgtcgacGGTAGTAATGTTACGTTCATTCAG |
| Nse1 Q18A M21A | JP891 | fw | GACAAGCATAAATTCATTCTTgcATATATAgcGTGTCGCACAGCTGGTGTTG |
| | JP892 | rev | CAACACCAGCTGTGCGACACgcTATATATgcAAGAATGAATTTATGCTTGTC |
| Nse4 L62C | oLV291 | fw | GAAGCAACCTTAGATGCTTTACTGtgTACTAAAACGGTTGATCTGGCTTC |
| | oLV292 | rev | GAAGCCAGATCAACCGTTTTAGTAcaCAGTAAAGCATCTAAGGTTGCTTC |
| Nse4 T63R | oLV612 | fw | AACCTTAGATGCTTTACTGCTTAgaAAAACGGTTGATCTGGCT |
| | oLV613 | rev | AGCCAGATCAACCGTTTTtcTAAGCAGTAAAGCATCTAAGGTT |
| Nse4 K64C | oLV295 | fw | GCAACCTTAGATGCTTTACTGCTTACTtgcACGGTTGATCTGGCTTCCA |
| | oLV296 | rev | TGGAAGCCAGATCAACCGTgcaAGTAAGCAGTAAAGCATCTAAGGTTGC |
| Nse4 T65R | oLV614 | fw | AGATGCTTTACTGCTTACTAAAAgGGTTGATCTGGCTTCCA |
| | oLV615 | rev | TGGAAGCCAGATCAACCcTTTTAGTAAGCAGTAAAGCATCT |
| Nse4 V66C | oLV417 | fw | GATGCTTTACTGCTTACTAAAACGtgTGATCTGGCTTCCATTAAAGC |
| | oLV418 | rev | GCTTTAATGGAAGCCAGATCAcaCGTTTTAGTAAGCAGTAAAGCATC |
| Nse4 D67C | oLV297 | fw | ATGCTTTACTGCTTACTAAAACGGTTtgTCTGGCTTCCATTAAAGCTAGG |
| | oLV298 | rev | CCTAGCTTTAATGGAAGCCAGAcaAACCGTTTTAGTAAGCAGTAAAGCAT |
| Nse4 L68C | oLV377 | fw | ACTGCTTACTAAAACGGTTGATtgcGCTTCCATTAAAGCTAGGCA |
| | oLV378 | rev | TGCCTAGCTTTAATGGAAGCgcaATCAACCGTTTTAGTAAGCAGT |
| Nse4 L62C T65R | oLV676 | fw | AGCAACCTTAGATGCTTTACTGtgTACTAAAAgGGTTGATCTGGCTTCCA |
| | oLV677 | rev | TGGAAGCCAGATCAACCcTTTTAGTAcaCAGTAAAGCATCTAAGGTTGCT |
| Nse4 del87-91 | oLV654 | fw | TTGGAAGGCCCAAGTTTAATATTGAAATTAAGCAATTCCTCAACTATCC |
| | oLV655 | rev | GGATAGTTGAGGAATTGCTTAATTTCAATATTAAACTTGGGCCTTCCAA |
| Smc5 E995Q | oLV494 | fw | GCTCCGTTTCGAATAGTTGATcAAATAAATCAAGGAATGGATCCTC |
| | oLV495 | rev | GAGGATCCATTCCTTGATTTATTTgATCAACTATTCGAAACGGAGC |
| Smc5 K57I | oLV674 | fw | TTTGATTATCGGTCCAAATGGGACAGGTAtaAGCACAATTGTTTCAG |
| | oLV675 | rev | CTGAAACAATTGTGCTTaTACCTGTCCCATTTGGACCGATAATCAAA |

| | | | |
|---|---|---|---|
| Smc6 L964A | oLV179 | fw | GCTAGACTCACGCAAgcATTGCAAGCTTTAGAAGA |
| | oLV180 | rev | TCTTCTAAAGCTTGCAATgcTTGCGTGAGTCTAGC |
| Smc6 L965A | oLV550 | fw | TGCTAGACTCACGCAACTAgcGCAAGCTTTAGAAGAGACGT |
| | oLV551 | rev | ACGTCTCTTCTAAAGCTTGCgcTAGTTGCGTGAGTCTAGCA |
| Smc6 Q966A | oLV552 | fw | CTAGACTCACGCAACTATTGgcAGCTTTAGAAGAGACGTTAC |
| | oLV553 | rev | GTAACGTCTCTTCTAAAGCTgcCAATAGTTGCGTGAGTCTAG |
| Smc6 A967C | oLV315 | fw | AGACTCACGCAACTATTGCAAtgTTTAGAAGAGACGTTACGAAGGC |
| | oLV316 | rev | GCCTTCGTAACGTCTCTTCTAAAcaTTGCAATAGTTGCGTGAGTCT |
| Smc6 L968A | oLV183 | fw | GCAACTATTGCAAGCTgcAGAAGAGACGTTACGAAG |
| | oLV184 | rev | CTTCGTAACGTCTCTTCTgcAGCTTGCAATAGTTGC |
| Smc6 E969A | oLV554 | fw | ACGCAACTATTGCAAGCTTTAgcAGAGACGTTACGAAGGCGT |
| | oLV555 | rev | ACGCCTTCGTAACGTCTCTgcTAAAGCTTGCAATAGTTGCGT |
| Smc6 E970A | oLV187 | fw | CTATTGCAAGCTTTAGAAGcGACGTTACGAAGGCGT |
| | oLV188 | rev | ACGCCTTCGTAACGTCgCTTCTAAAGCTTGCAATAG |
| Smc6 T971A | oLV556 | fw | ACTATTGCAAGCTTTAGAAGAGgCGTTACGAAGGCGTAATG |
| | oLV557 | rev | CATTACGCCTTCGTAACGcCTCTTCTAAAGCTTGCAATAGT |
| Smc6 L972A | oLV558 | fw | TGCAAGCTTTAGAAGAGACGgcACGAAGGCGTAATGAAATGT |
| | oLV559 | rev | ACATTTCATTACGCCTTCGTgcCGTCTCTTCTAAAGCTTGCA |
| Smc6 R973A | oLV189 | fw | GCTTTAGAAGAGACGTTAgcAAGGCGTAATGAAATGTG |
| | oLV190 | rev | CACATTTCATTACGCCTTgcTAACGTCTCTTCTAAAGC |
| Smc6 R974A | oLV191 | fw | CTTTAGAAGAGACGTTACGAgcGCGTAATGAAATGTGGAC |
| | oLV192 | rev | GTCCACATTTCATTACGCgcTCGTAACGTCTCTTCTAAAG |
| Smc6 R975A | oLV193 | fw | GAAGAGACGTTACGAAGGgcTAATGAAATGTGGACCAAATTTC |
| | oLV194 | rev | GAAATTTGGTCCACATTTCATTAgcCCTTCGTAACGTCTCTTC |
| Smc6 S1045R | oLV498 | fw | AAGTCAGCGTTCAAGGATTAcgAGGGGGTGAAAAATCTTTTG |
| | oLV499 | rev | CAAAAGATTTTTCACCCCCTcgTAATCCTTGAACGCTGACTT |
